# The precision of attention controls attraction of population receptive fields

**DOI:** 10.1101/2025.06.16.659874

**Authors:** Sumiya Sheikh Abdirashid, Tomas Knapen, Serge O. Dumoulin

**Affiliations:** Spinoza Centre for Neuroimaging, Amsterdam, Netherlands; Computational Cognitive Neuroscience and Neuroimaging, Netherlands Institute for Neuroscience, Royal Netherlands Academy of Arts and Sciences, Amsterdam, Netherlands; Experimental and Applied Psychology, Vrije University Amsterdam, Amsterdam, Netherlands; Experimental Psychology, Helmholtz Institute, Utrecht University, Utrecht, Netherlands

**Keywords:** retinotopy, visual attention, population receptive fields, 7T MRI, computational modeling

## Abstract

We alter our sampling of visual space not only by where we direct our gaze but also by where and how we direct our attention. Attention attracts receptive fields toward the attended position, but our understanding of this process is limited. Here we show that the degree of this attraction towards the attended locus is dictated not just by the attended position, but also by the precision of attention. We manipulated attentional precision while using 7T fMRI to measure population receptive field (pRF) properties. Participants performed the same color-proportion detection task either focused at fixation (0.1° radius) or distributed across the entire display (more than 5° radius). We observed BOLD response amplitude increases as a function of the task, with selective increases in foveal pRFs for the focused attention task and vice versa for the distributed attention task. Furthermore, cortical spatial tuning changed as a function of attentional precision. Specifically, focused attention more strongly attracted pRFs towards the attended locus compared to distributed attention. This attraction also depended on the degree of overlap between a pRF and the attention field. A Gaussian attention field model with an offset on the attention field explained our results. Together, our observations indicate the spatial distribution of attention dictates the degree of its resampling of visual space.

## Introduction

Attention is often described as a zoom lens because it enhances visual perception at the attended location (Carrasco, 2011; Eriksen & St James, 1986). Voluntary spatial attention alters neuronal activity and representations of visual space (Carrasco, 2011; Datta & DeYoe, 2009; Fischer & Whitney, 2009; Kastner et al., 1999; Kay et al., 2015, 2015; Liu et al., 2022; McAdams & Maunsell, 1999; Moran & Desimone, 1985; Tootell et al., 1998; Tünçok et al., 2024; Vo et al., 2017). Attention attracts receptive fields toward the attended location, resulting in better sampling at the attended location (Klein et al., 2014, 2018; van Es et al., 2018; Womelsdorf et al., 2006, 2008). This receptive field attraction is thought to underlie improved spatial resolution and enhanced perception at the attended location – akin to a neural implementation of a zoom lens (Anton-Erxleben & Carrasco, 2013; Baruch & Yeshurun, 2014; Klein et al., 2016). However, the factors that determine the degree of attraction towards the attended locus (i.e., the magnification of the ‘zoom lens) has yet to be elucidated.

Attention field models are widely used to describe and predict the influence of attention on cortical visual responses (Klein et al., 2014; Puckett & DeYoe, 2015; Reynolds & Heeger, 2009; Womelsdorf et al., 2008). The Gaussian attention field model summarizes both attention and receptive fields as Gaussians and their interaction as a multiplication of Gaussians (Klein et al., 2014; Reynolds & Heeger, 2009; Womelsdorf et al., 2008) (Equation 1, Figure 1A). This model predicts that the precision of the attention field determines the attraction of receptive fields (Klein et al., 2014; van Es et al., 2018; Womelsdorf et al., 2006). In line with these predictions, attentional precision alters response gain and contrast gain (Herrmann et al., 2010; Itthipuripat et al., 2014). Specifically, the attention field model predicts that more precise, focused attention (i.e. a smaller Gaussian standard deviation) will result in stronger reweighting towards the attended locus (Figure 1A), under conditions of equal task difficulty and performance.

**Figure 1.**
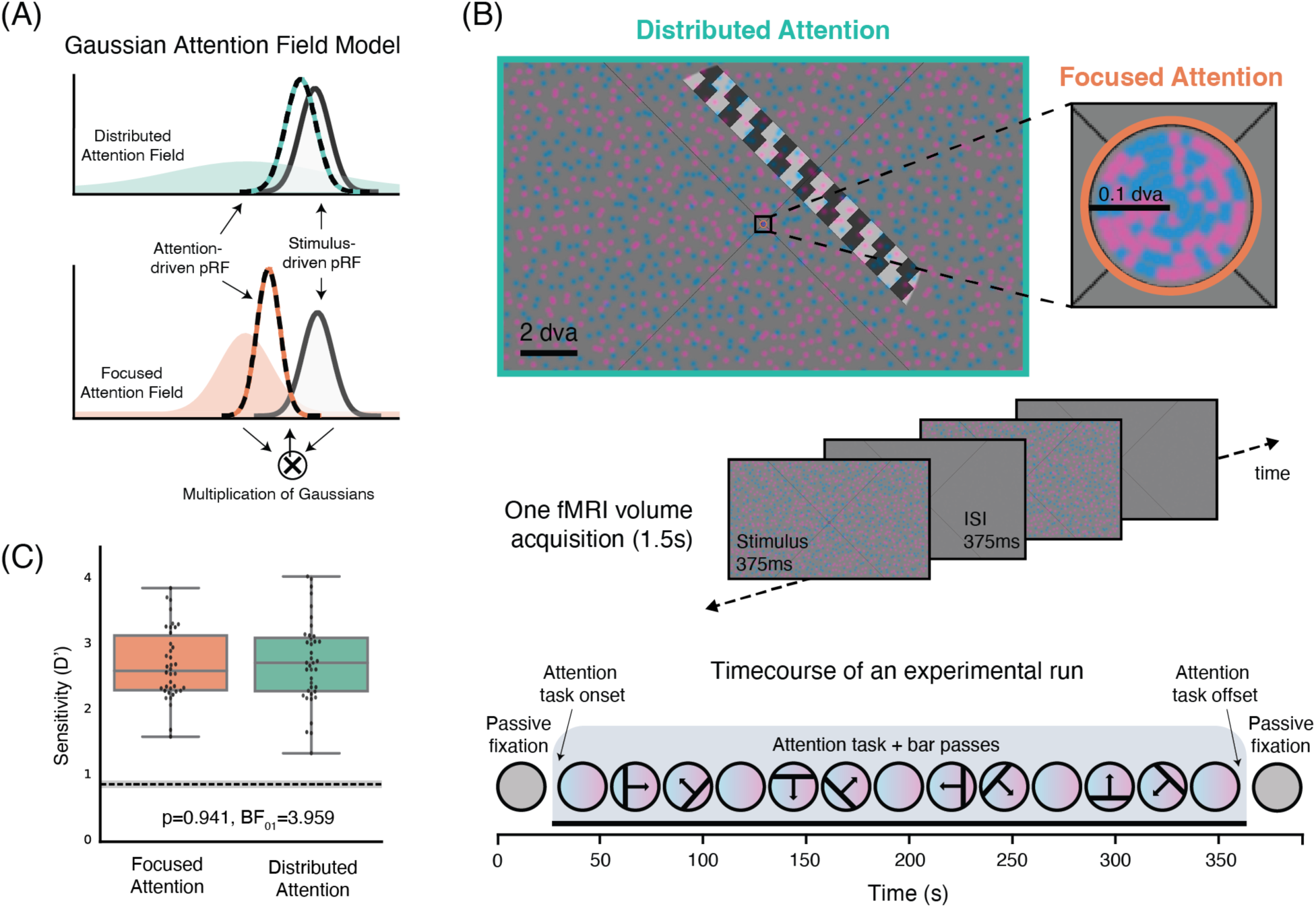
**a)** The attention field model describes the influence of attention as a multiplication of a Gaussian attention field with a Gaussian population receptive field (Klein et al., 2014; Reynolds & Heeger, 2009; van Es et al., 2018; Womelsdorf et al., 2008). Focused attention (a narrower Gaussian) attracts resulting attention-driven receptive fields more compared to distributed attention (a wider Gaussian). **b)** Top panel: one stimulus example. A color-proportion discrimination task is performed either across the entirety of the screen (distributed attention) or within the fixation circle (focused attention). Middle panel: illustration of the stimulus sequence during 1 fMRI volume acquisition. Bottom panel: time course of one experimental run. Each run begins and ends with a mean-luminance fixation block, after which the attention task and bar passes begin. **c)** Sensitivity (D’) for the focused and distributed attention conditions are indistinguishable.

Our aim was to investigate whether precision of attention indeed dictates the degree of attraction of receptive fields towards the attended locus. We developed a paradigm to reconstruct population receptive field properties (Dumoulin & Wandell, 2008) while we manipulated the precision of attention (Figure 1B, Supplementary Movie 1). To maximize the difference in attentional precision, we used two conditions: attention directed at fixation (0.1° radius) and attention distributed across the entire screen (∼ 5° x 11°).

We observed BOLD response amplitude differences as a function of the task. pRFs falling within the stimulus extent of the focused task, showed increased BOLD amplitude for the focused task. Similarly, pRFs falling within the stimulus extent of the distributed task showed increased BOLD amplitude for the distributed task. We also observed that the precision of attention alters population receptive field properties within visual field maps and along the visual hierarchy. The influence of attentional precision varies with increasing distance from the attentional locus. Additionally, these observations were well captured by a Gaussian attention field model with an offset on the attention field.

## Materials & Methods

### Participants

In the final analysis, 10 participants were included (five female, aged 24-42); 3 participants were excluded (one for not completing all scanning sessions, one due to poor signal-to-noise ratio because of distance from the coil, and one because behavioral performance in the two conditions not being matched). All participants had normal or corrected-to-normal visual acuity. This study was approved by the Human Ethics Committee of Vrije Universiteit Amsterdam and was conducted with informed written consent of all participants.

### Experimental design

Visual stimuli were presented on an MR compatible Cambridge Research Systems LCD screen (69.84 x 39.29 cm, 1920 × 1080 pixels, 120-Hz refresh rate). The screen was located outside of the bore and participants viewed it through a mirror mounted on the coil, the screen was 210 cm from the mirror. For training on the task outside of the scanner, we used a comparable setup with an identical screen, screen distance, and visual stimuli.

The visual stimuli were created using the PsychoPy python package (Peirce, 2007) and the exptools2 wrapper for PsychoPy (https://github.com/VU-Cog-Sci/exptools2). A visual field mapping experimental design was used in combination with a demanding attentional task. A black diagonal fixation cross filled the screen and a fixation circle (0.1° visual angle radius) was placed in the center of the screen. An overview of the stimulus and task can be found in Figure 1B and Supplementary Movie 1.

Participants were instructed to fixate, to conduct a task, and ignore the checkerboard bar traversing the screen. The task was either focused within the fixation circle (0.1° visual angle radius) or distributed across the entire screen (5.6° x 11.1° visual angle). in both the focused and distributed conditions, the visual stimulus presented and the task participants carried out were identical. The only difference between these two conditions was the spatial extent to which participants would need to focus or distribute their attention in order to do the task.

Across the screen (0.5°-5° height, 0.5°-11° width) a number of pink (RGBA:1,0.35,0.87,0.25) and blue (RGBA: 0.1,0.6,1,0.25) dots were presented on a mean luminance background. Simultaneously, within the 0.1° radius fixation circle, a number of identically colored dots were presented on a mean luminance background. Each trial consisted of 375ms color-dot stimulus presentation and 375ms interstimulus interval. The dot positions jittered trial-by-trial and each dot color was updated trial-by-trial.

The task was a color-proportion detection task. Participants were instructed to detect target trials where the proportion of dots filling the screen deviated from 50-50, becoming overall more pink or more blue. On non-target trials (i.e. no-response trials), the proportion of pink:blue dots was 50-50. Roughly 1 in 12 stimulus presentations was a target, a jitter was used to avoid predictability. This proportion detection task becomes more difficult the closer the target trials are to the 50-50 proportion. To titrate the difficulty of the task and to match performance across conditions and participants, the target proportion was adjusted. Target proportions were set independently for each condition. A reaction time window 1s was used to log correct responses.

At the beginning of each run, participants were assigned a task condition (focused at fixation or distributed across the whole screen) that spanned the duration of the run. There were two mean-luminance fixation blocks at the start (20s) and end (30s) of every run, where no color-dot stimulus was presented. After the first 20s fixation block the color-dot stimulus was presented and the attentional task began.

The visual field mapper started 15s after the start of the color-dot stimulus. A checkerboard bar (1.25° width, 50% Michelson contrast) was used as a visual field mapper. It traversed the screen within a circular aperture with a diameter of ∼10° of visual angle. The bar swept the screen with four orientations (0°, 45°, 90°, 135°) and four motion directions (perpendicular to bar orientation), resulting in a total of eight bar passes per run. The checkerboard pattern within the bar also moved parallel to bar orientation. The bar motion was time-locked to the MRI acquisition and moved 0.625° each TR (1.5s). There was a 15s gap after every two bar sweeps, during which the attentional task continued. The bar sweeps were only presented within the duration of an attentional task. After the last bar sweep there is an additional 30s of the task followed by 30s of mean luminance fixation.

All participants were trained on the task prior to scanning to avoid learning effects during scanning. During training we also used a 2-alternative forced-choice (2AFC) version of the task to set an initial task difficulty. In this task, on each trial participants would respond whether the stimulus presented was more pink or more blue. Again, participants either focused attention within the fixation circle or distributed attention across the entire screen. This task was used to obtain a psychometric curve for each participant for the two attention conditions. These psychometric curves were used to choose values of task difficulty that were matched for the two conditions and matched across subjects. Following this 2AFC task, participants trained on the color-proportion detection task (described above) to be used in the scanner.

For each participant, 6-8 runs of this experiment were collected in the scanner as well as an independent pRF mapping session.

### Independent visual field mapper

In addition to this experimental paradigm, a standard visual field mapping session was collected for all participants. The visual stimulus and task follow those outlined in the methods of (Aqil et al., 2021; Dumoulin & Wandell, 2008). This visual field mapper used a contrast-defined checkboard bar traversing the screen on a mean luminance background. It moved in 8-motion directions (4 cardinal, 4 oblique). Participants were instructed to fix their gaze on a dot at the center of the screen and press a button whenever the dot changed color. Attention towards the fixation may therefore influence pRF properties in the mapper, however, the independent mapper can never be an estimate of attention-free pRF properties as attention is always employed even without task demands. The primary use of the visual field mapper was to provide independent estimates and prevent circular analyses and regression to the mean (Stoll et al., 2022). The pRF parameters acquired from this independent mapper session were used for all the binning along eccentricity (Figure 2B-C, Figure 3D-E).

**Figure 2:**
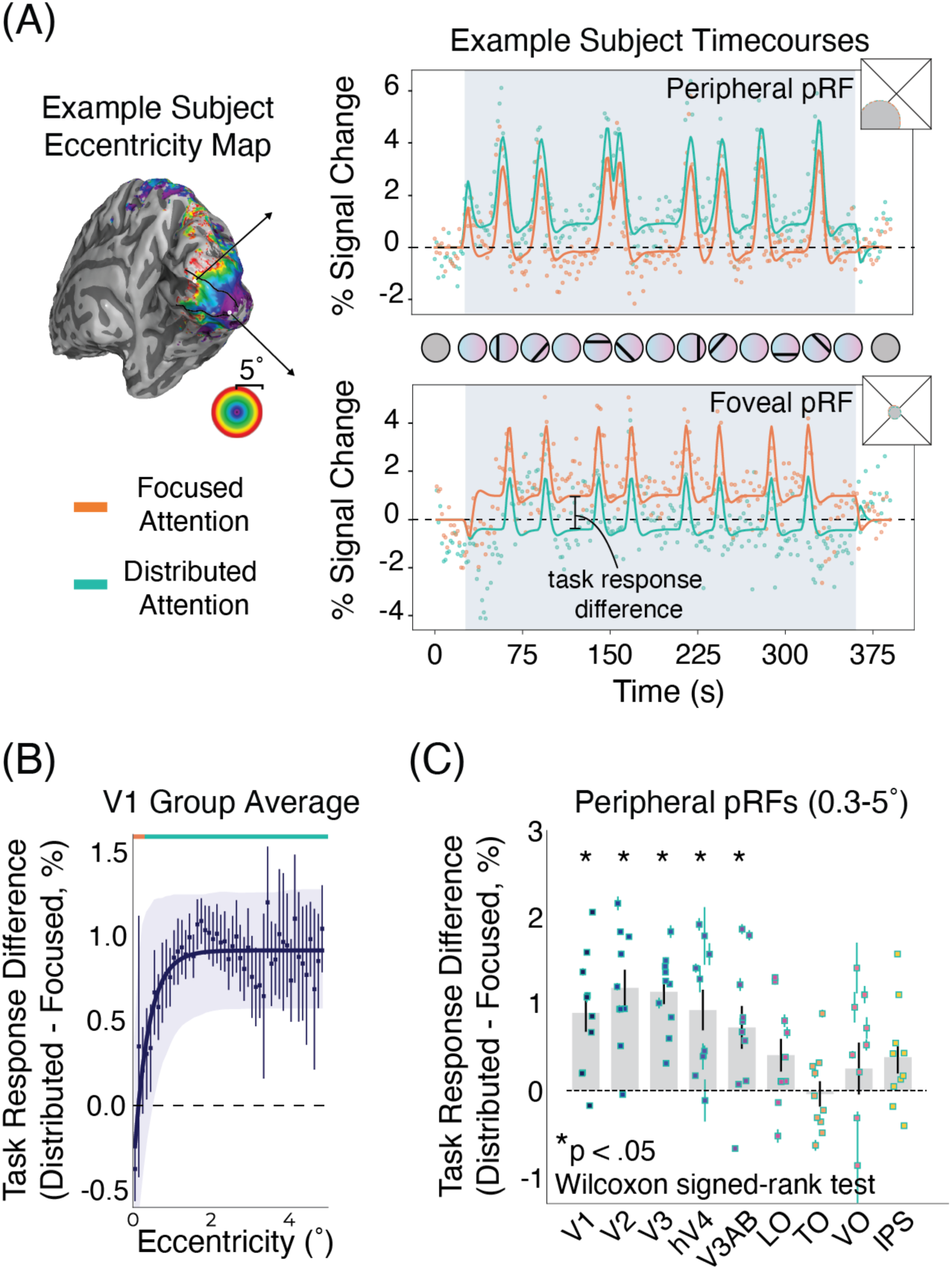
Increased BOLD task responses are dependent on eccentricity and attentional precision. **a)** Example subject V1 peripheral pRF timecourse (top) and V1 foveal pRF timecourse (bottom). showing response differences for the distributed and focused attention task. For the peripheral pRF, task response is increased for the distributed task relative to the focused task. Conversely, for the foveal pRF task response is increased for the focused task relative to the distributed task. **b)** Task response difference (distributed task response minus focused task response) for pRFs in V1, binned for both tasks based on an independent pRF condition. The orange line at the top of the plot denotes the stimulus extent of the focused attention task, teal line denotes the stimulus extent of the distributed attention task. The results show that the response difference varies across eccentricity in line with the task extent. Each individual point is an average of all participants within each eccentricity bin. The solid purple line is an exponential function fit to the data. The shaded area denotes a bootstrapped 95% confidence interval of the exponential curve fit. **c)** Task response difference across the visual hierarchy for pRFs within the distributed task stimulus extent (0.3-5 dva). Each point is an individual participant average. Error bars are SEM. These results suggest that the response amplitude elicited by the color-dot stimulus is modulated by eccentricity and task.

**Figure 3:**
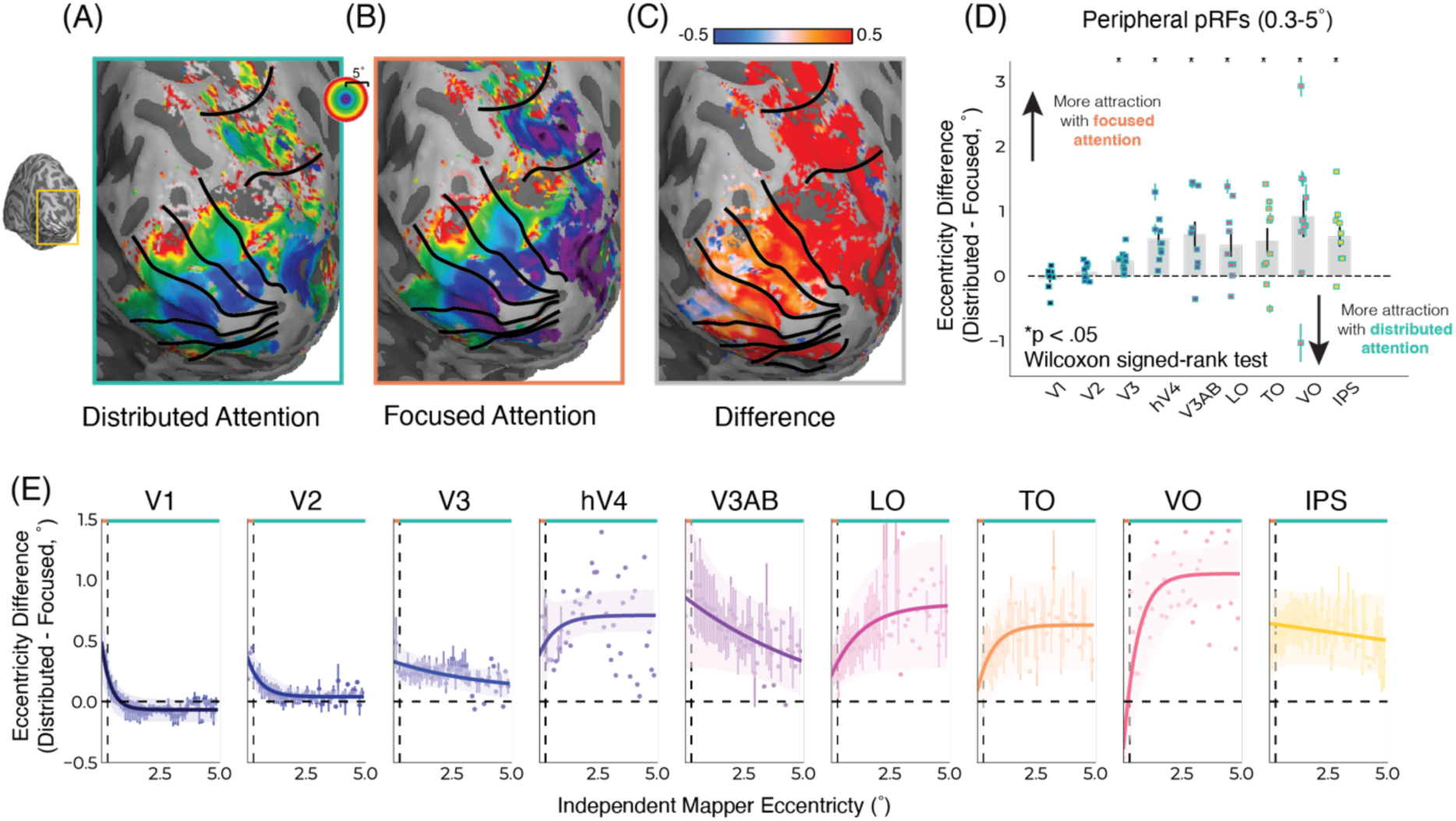
Focused attention more strongly attracts pRF position towards the fovea. **a)** One example subject eccentricity map for distributed attention and **b)** focused attention for one example subject. **c)** Difference in eccentricity maps (distributed minus focused). Vertices were masked based on variance explained (R^2^ ≥ 0.1) and eccentricity (< 5°). **d)** Eccentricity difference (distributed minus focused) across the visual hierarchy for pRFs falling within the distributed color task stimulus (0.3-5° radius). Values per participant are shown in the scatter plot nested within each bar, group averages in bar plots, and error bars are SEM. **e)** Eccentricity difference (distributed minus focused) for pRFs in each visual area binned for both tasks based on an independent pRF condition. The orange line at the top of the plot denotes the stimulus extent of the focused attention task (0-0.3° radius), the teal line denotes the stimulus extent of the distributed attention task (0.3-5° radius), and the dashed vertical line the boundary between the two tasks. The results show that pRF attraction is not uniform across eccentricity, with pRFs in V1 and V2 closest to the locus of attention showing more attraction (positive eccentricity difference) that is not visible in pRFs further away. Each individual point is an average of all participants within each eccentricity bin. The thick solid lines are exponential functions fit to the data. The shaded area denotes a bootstrapped 95% confidence interval of the exponential curve fit. All error bars are SEM.

### Statistics

We used both Bayesian and frequentist statistics to determine whether the two attentional conditions were matched in behavioral performance. The discriminability index (D’) for each condition was compared across subjects (Figure 1C). JASP Bayesian ANOVA was performed to test for evidence in favor of the null hypothesis. D prime was the dependent variable, the attention conditions were the fixed factors and subject and run were modeled as random effects variables (van Doorn et al., 2021). We also used a frequentist linear mixed model, with the same dependent variable, fixed effect variables, and random effect variables.

For the fMRI data, first a Shapiro-Wilk test was used to test for normality, the data was not normally distributed, so we used non-parametric statistical tests for subsequent analyses. We performed a Wilcoxon signed-rank test to determine whether task response (Figure 2C) and eccentricity (Figure 3D) differed significantly between conditions. To correct for multiple comparisons the false-discovery rate (FDR) correction was used.

### Eye-tracking

Prior to carrying out the experiment in the MRI participants were trained on an equivalent set-up (same screen properties and distance). During this training gaze position was recorded using an Eyelink 1000 (SR Research, Osgoode, Ontario, Canada). The eye-tracker was calibrated using a 5-point procedure. Due to technical constraints eye-tracking data could not be recorded for all subjects during the scanning session. For these subjects without eye-tracking data during scanning, gaze position recorded during the training session was used.

### MRI data acquisition

All scans were acquired on a Philips Achieva 7T scanner with a 32-channel Nova Medical head coil.

T1-weighted (T1w) MP2RAGE and T2-weighted (T2w) turbo-spin echo structural MRI scans were acquired at a resolution of 0.7mm isotropic (T1w: FOV = 220 x 220 x 200 mm^3^, matrix = 352 x 352 x 263, TR / TE = 6.2 ms / 3 ms, flip angle_1_ / flip angle_2_ = 5° / 7°, TR_MP2RAGE_/TI_1_/TI = 5500 ms/ 800 ms/2700 ms, duration = 9 min 45 s; T2w: FOV = 245 x 245 x 184 mm^3^, matrix = 352 x 349 x 263, TR/TE = 3000/390 ms, TSE-factor = 182, duration = 7 min).

For functional scans, a two-dimensional EPI sequence was used with 60 coronal slices, 216×216 FOV, 1500ms TR, 23ms TE, 65° flip angle, PA phase encoding direction, and 1.7mm isotropic resolution. To avoid start-up magnetization transients, the first 10s of recorded data were discarded. 4 top-up scans with opposing phase-encoding directions were collected throughout each scan session, which was later used for susceptibility distortion correction. Each run had a scan duration of 330s (independent pRF mapper) or 390s (attentional task experiment).

### Preprocessing

The anatomical preprocessing pipeline by Heij and colleagues (Heij et al., 2023) was used (https://github.com/gjheij/linescanning). MP2RAGE images were combined from the first and second inversions. Anatomical surfaces were reconstructed using Freesurfer 7.2 combining T1 and T2 images. Functional data was sampled to the reconstructed surface using the default fMRIprep pipeline (version 21; Esteban et al., 2019). BOLD volumes were averaged, linear de-trended, and converted to percent signal change prior to fitting procedures. This process was carried out separately for each experimental condition and separately for each participant.

### Attention field models

The attention field (AF) model is given by Equation 1:

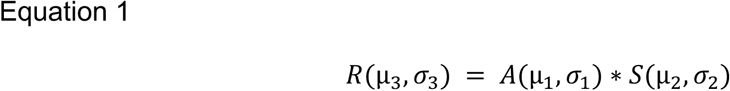

where µ,σ are the position and size of a Gaussian receptive field, A represents the Gaussian attention field and S represents the Gaussian stimulus-driven receptive field. R is the resulting attention driven receptive field, resulting from the multiplication of the stimulus-driven receptive field and Gaussian the attention field. This model (Klein et al., 2014) has an analytical solution for the attention-induced position changes (Equation 2):

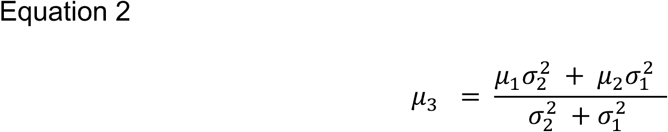

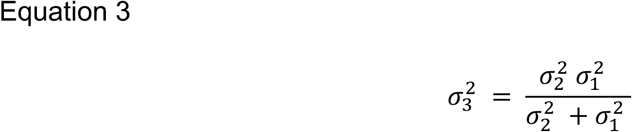

The position of the attention-driven receptive field is estimated by μ_3_. μ_1_ and σ_1_ are the position and size of the Gaussian attention field respectively, while μ_2_ and σ_2_ are the position and size of the Gaussian stimulus-driven receptive field.

The AF+ model (Reynolds & Heeger, 2009; Womelsdorf et al., 2008), adds an offset to the attention field (Figure 4) and is given by Equation 3:

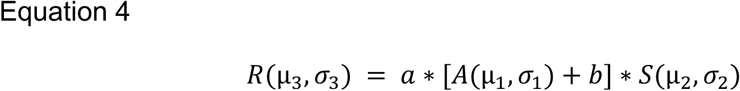

**Figure 4:**
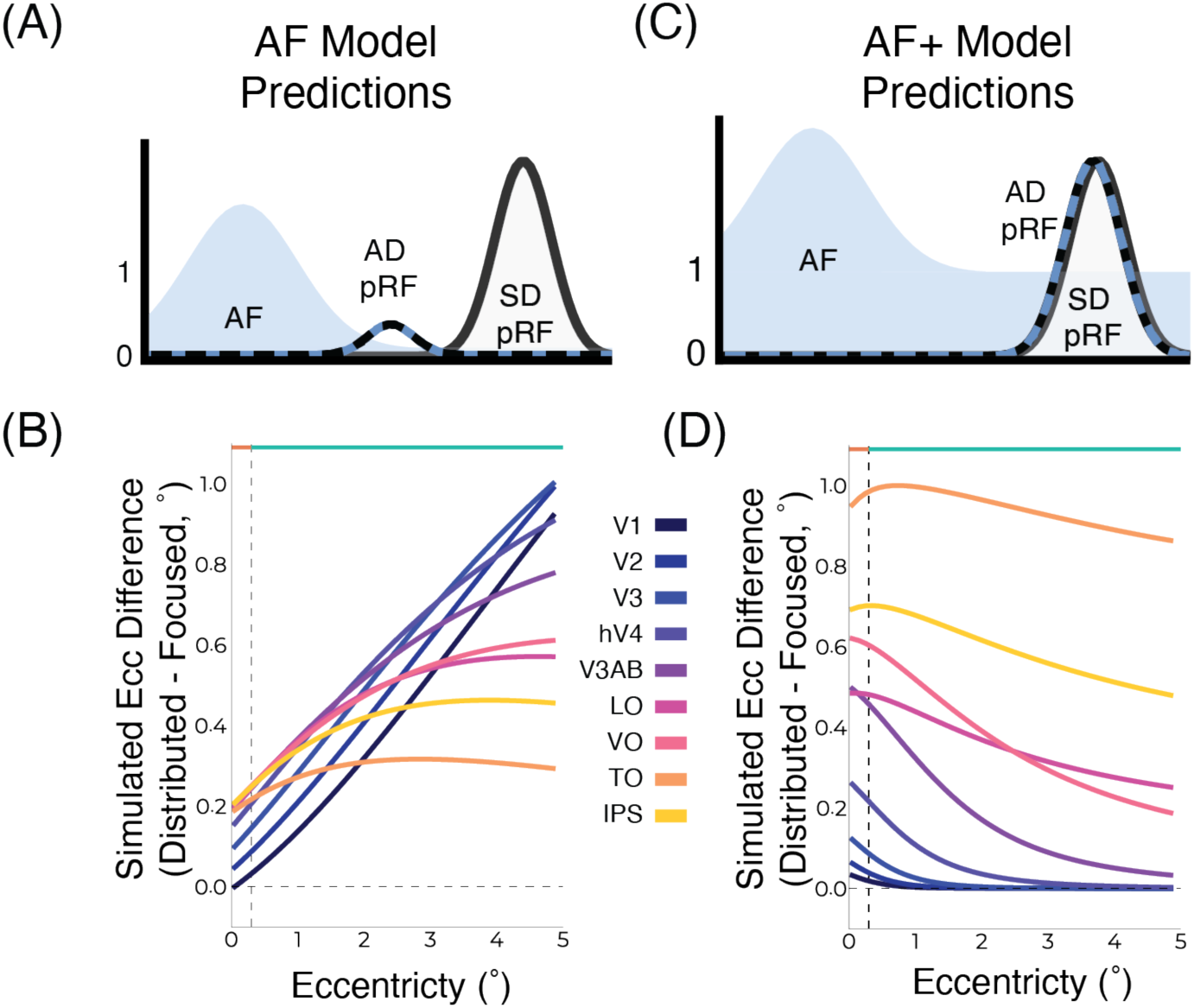
Adding an offset to the attention field improves predictions.: **a)** Predictions of the Gaussian attention field model without an offset (AF model; Equation 1) when the stimulus-driven pRF (SD-pRF) and attention field (AF) are far apart the resulting Gaussian has a very small amplitude. **b)** Simulated predictions of this AF model, showing increasing or plateauing eccentricity difference with increasing eccentricity. **c)** The attention field model with an offset on the attention field (AF+ model; Equation 3) does not result in infinitely small amplitudes when SD-pRF and AF are far apart, instead the relative influence of the attention field becomes negligible and the resulting Gaussian is the SD-pRF. Instead when the SD-pRF and AF are closer, the influence of the attention field is larger and there is a greater position change. **D.** Simulated predictions of the AF+ model. For V1-3 there are only position differences foveally and peripherally there are no differences. In higher visual areas there is a plateau of differences along the eccentricity range. This model better captures the patterns in the data.

This AF+ model has two additional parameters, *b,* which is a constant added to the attention field and *a,* a scaling factor.

### pRF and GLM fitting

In every run, a block design was nested into the event-related experiment design. The blocks were: 14 TRs (21s) passive fixation, 224 TRs (5.6 minutes) attention task with pRF mapping, and 21 TRs (31.5s) passive fixation (Figure 1B, bottom panel). We first truncated the timecoures to exclude the passive fixation blocks and any transient responses at the onset and offset of the task. pRF fitting was then performed on these truncated timecourses using the Python package Prfpy (Aqil & Knapen, 2023) (https://github.com/VU-Cog-Sci/prfpy), following the procedure outlined in Dumoulin and Wandell (2008). Only pRFs with a variance explained of 0.1 or greater and with an eccentricity less than 5° were used for subsequent analyses.

Next, on the uncut timecourses, a GLM was used to account for the blocks in the experimental design. The two passive fixation blocks served to estimate a BOLD baseline outside of the attentional task. The predictors in the design matrix were: the predicted time course resulting from the pRF fitting, the attentional task, and two nuisance regressors at the TR for the task onset and offset. All regressors were convolved with a canonical hemodynamic response function. Least squares minimization was used. The beta value for the attentional task obtained from the GLM is subsequently referred to as the “task response” (Figure 2).

### Attention field model simulations

Using the size-eccentricity curves from the independent pRF mapper, pRFs were simulated to predict how their positions would be altered by simulated attention fields. To do this, we used the attention field model equations (above). Since stimulus-driven pRFs are only hypothetical and can not be measured we simulated pRF positions and sizes to use as the stimulus-driven input. Two simulated attention fields were used to mimic our experimental design: one with a small standard deviation and one with a larger standard deviation. Both simulated attention fields were centered at 0. These simulated attention fields when multiplied by the simulated stimulus-driven pRFs resulted in position differences. These position differences were used to fit the simulated attention fields to the observed data, the eccentricity differences shown in Figure 3E. We used the exponential functions which captured the observed position changes across the eccentricity range (Figure 3E). These simulated position differences are shown in Figure 4 B and D. Klein et al. (2014) demonstrated that a single attention field could be used to explain effects along the entire visual hierarchy. Correspondingly, we used one shared attention field across the entire visual hierarchy within each of our conditions. We further investigated two formulations of the attention field model: the attention field model with an offset on the attention field (AF+) and without an offset on the attention field (AF) (Figure 4). For the AF+, there is no formulaic solution. As a result, for each simulated attention driven pRF arising from a multiplication with the AF+, we fit a new Gaussian function to provide an estimate for the center of mass of the resulting distribution.

### Regions of interest (ROIs)

Visual field maps of eccentricity, polar angle, and pRF size were obtained from the independent pRF mapper for each participant. These maps were used to draw regions of interest (V1, V2, V3, hV4, V3ab, LO, TO, VO, Lower IPS, Upper IPS) (Amano et al., 2009; Swisher et al., 2007; Wandell et al., 2005, 2007).

## Results

### Behavioral performance for distributed and focused spatial attention is matched

As intended, participants performed the task comparably in the two conditions (Fig 1C) with an average *D’* and standard error of 2.709 ± 0.089 and 2.745 ± 0.103 for the focused and distributed attention conditions, respectively. A permutation test was performed to obtain a null distribution for *D’* (Fig 1C). All participants’ performance surpassed that of the null distribution, with no differences between the two attention conditions (linear mixed model, *p*=0.941, i.e., no significant difference; Bayesian *ANOVA BF_01_* 3.959, i.e., strong evidence that performance was similar). We also observed no significant differences in reaction time or criterion. Furthermore, all subjects were trained to fixate comparably for the two conditions. Due to technical constraints we could not record eye-tracking for all subjects in the scanner, however, we collected this data outside of the scanner for all subjects. Fixation was comparable for our two experimental conditions. Thus, the focused and distributed attention conditions were matched in terms of behavior.

### Responses elicited by color-dot stimuli are modulated by eccentricity and attentional precision

Attention increases responses in the attended parts of the visual field in line with both fMRI studies (Datta & DeYoe, 2009; Kastner et al., 1999; Müller et al., 2003; Puckett & DeYoe, 2015; Tootell et al., 1998) and neurophysiology studies (Connor et al., 1996; Luck et al., 1997; Moran & Desimone, 1985; Reynolds et al., 2000). For V1 pRFs falling within the extent of the focused attention task (0 - 0.3° radius), there was a higher BOLD task response for the focused attention task compared to the distributed attention task (Fig 2A, B). Conversely, for V1 pRFs falling within the extent of the distributed attention task (0.3 - 5° radius), BOLD task response was higher for the distributed attention task compared to the focused attention task (Fig 2A, B). This result demonstrates a selective enhancement of the BOLD response for V1 pRFs encoding parts of the visual field where the task is being performed.

A limited number of pRFs fell within the 0.3° extent of the focused attention task, yet, this pattern in the BOLD task response can still be observed across the visual hierarchy in peripheral pRFs. Across V1-3, hV4 and V3A/B, peripheral pRFs (0.3 - 5° radius) showed positive task response differences (i.e. higher BOLD task response for the distributed task compared to the focused task) reflecting those seen in V1 (Fig 2C; Wilcoxon signed-rank test, FDR corrected *p_V1_*=0.009, *p_V2_*=0.009, *p_V3_*=0.009, *p_hV4_*=0.009, *p_V3A/B_*=0.025). In some visual areas (LO, TO, VO, IPS) this distinction was not significant. This could be due to the increased pRF sizes and smaller visual field maps in higher visual areas resulting in less clear distinction of foveal and peripheral signals.

### Focused attention more strongly attracts pRF position preference

Across the visual hierarchy, precision of attention alters changes in visual spatial tuning. Eccentricity maps for the focused and distributed attention conditions are shown for one example subject in Figure 3A and B respectively. Visibly, there are position differences in these maps with more foveal pRF positions for the focused attention task compared to the distributed attention task. Vertex-wise subtracting the pRF positions from each condition reveals smaller eccentricity values in the focused condition compared to the distributed condition. This means more foveal pRFs for the focused attention condition, indicating stronger attraction towards the attended locus (Figure 3C). Notably, this pattern of position changes can not be explained by foveal-peripheral differences resulting from the task. Were this the case, we would expect the opposite pattern of results, increased attentional effects for the distributed, more peripheral, task (Anton-Erxleben & Carrasco, 2013; Carrasco, 2011; Rosenholtz, 2016). Instead, we see the opposite, stronger attraction towards the attended locus in the focused condition.

These results are in line with predictions from the attention field model (Figure 1A). Moreover, these differences in attraction between focused and distributed attention are more pronounced higher up the visual hierarchy, plotted for peripheral pRFs in Figure 3D. The attention field model also predicts this, as pRF sizes increase up the visual hierarchy so does the expected attraction towards the attended locus. The difference in position between the two tasks significantly differed for V3, hV4, V3A/B, LO, TO, VO, and IPS (Wilcoxon signed-rank test, FDR corrected *p_V3_*=0.009, *p_hV4_*=0.009, *p_LO_*=0.035, *p_TO_*=0.035, *p_VO_*=0.035, *p_IPS_*=0.012). While there were no significant differences in position change in V1 and V2 (Figure 3D), examining pRFs across eccentricity reveals non-linearities in the response depending on distance from the attended locus (Figure 3E).

### Attraction depends on distance from the attentional locus

In early visual cortex (Figure 3E, V1-3), greater attraction for focused attention compared to distributed attention was only seen in foveal pRFs (0-0.3° eccentricity) falling within the stimulus range of the focused attention task. However, for peripheral pRFs (0.3-5° eccentricity), falling within the eccentricity range of the distributed task, no significant differences in positions were observed. This result is better captured by an attention field model with an offset on the attention field (Figure 4). Without an offset, the attention field model predicts the attraction of a receptive field towards the attended locus should linearly increase with distance from the attended locus (Klein et al., 2014). Further up the visual hierarchy where pRFs are larger, a constant difference between focused and distributed attention conditions was observed, with narrow attention resulting in a stronger attraction along the entire eccentricity range (Figure 3E). These results indicate that attraction of pRFs towards the attended locus is determined not only by the precision of attention but also by the distance between a given receptive field and the attentional locus, with greater attentional influence resulting from greater overlap.

Importantly, this observed pattern of results can not be explained by foveal peripheral differences. Both tasks are centered at fixation, making the focused attention task more foveal and the distributed task more peripheral. Many examples in the literature point to larger attention effects in the periphery because spatial resolution is poorer there (Anton-Erxleben & Carrasco, 2013; Carrasco, 2011; Rosenholtz, 2016). If our observations were purely a result of foveal-peripheral differences, we would expect stronger attention effects in the distributed condition since it is more peripheral. What we observed was the opposite, a stronger attraction towards the attended locus during the focused (i.e. foveal) task. Furthermore, when looking along the eccentricity range, this effect *increased* as pRFs became more peripheral (Fig. 3E), in keeping with the literature (Anton-Erxleben & Carrasco, 2013; Carrasco, 2011; Klein et al., 2014; van Es et al., 2018). Thus, we can deduce the results are not due to foveal-peripheral differences.

We observed that the influence of attention is also a function of the amount of overlap between the stimulus-driven pRF (SD-pRF) and the attention field (AF). An attentional field model without an offset on the attention field predicts the position change increase or plateau with distance from the attended locus and with increasing pRF size (Figure 4B). This AF model does capture the pattern of position we see in higher visual areas (LO, TO, VO, IPS), however, it fails to capture the increase in position changes foveally in early visual areas (V1-V3; Figure 3E). Moreover, this model further predicts that as the SD-pRF and the AF become increasingly far apart, the resulting attention-driven pRF (AD-pRF) amplitude approaches zero (Figure 4A). For this reason, this model discards the amplitude parameters and only takes into account the Gaussians’ positions and sizes.

Adding an offset of parameter to the attention field (Figure 4C; Equation 2) (Reynolds & Heeger, 2009; Womelsdorf et al., 2008) better captures and contextualizes the results observed. With an offset on the attention field, the relative contribution of the attention field depends on the distance between the SD-pRF and the attentional locus. The influence of attention decreases with distance from the attended locus (Figure 4D). As the SD-pRF becomes farther from the attended locus, this model predicts that the resulting AD-pRF approaches the SD-pRF, i.e. less attraction towards the attended locus for pRFs that are farther away. This phenomenon is present in our data within the early visual cortex (Figure 3E, V1-3), characterized by smaller pRFs, resulting in less potential overlap with the AF. Indeed, as this model predicts, we observe that the influence of attention is present in pRFs close to the attentional locus declines with pRFs that are more distant from the attended locus.

## Discussion

We used an fMRI attentional experiment to selectively manipulate the precision of attention while clamping attended location, visual stimulus, and task difficulty. This design narrowly constrained the spatial extent of attention in one condition and maximally distributed it in another. In comparing these conditions, we saw that attentional precision influenced two separate components of cortical responses. First, we observed modulation of the BOLD baseline in response to the task. Specifically, increased BOLD task response in cortical regions which spatially encode color-dot stimulus. This result is in keeping with the attentional gain modulation literature (Brefczynski & DeYoe, 1999; Kastner et al., 1999; Maunsell, 2015; McAdams & Maunsell, 1999; Moran & Desimone, 1985; Treue & Trujillo, 1999). Second, we observed changes in cortical spatial tuning. Within an ROI, there was a greater overall re-weighting of the Gaussian population responses towards the attended locus for focused compared to distributed attention. This result follows predictions of the attention field model (Klein et al., 2014; Reynolds & Heeger, 2009; Womelsdorf et al., 2008) and demonstrates that precision of attention acts as a ‘magnification dial’ for the zoom lens of attention.

The attraction of pRFs towards the attended locus was not uniform across the eccentricity range. Our experimental design intentionally constrained the attention field in the focused attention condition, allowing us to probe how distance from the attention field influences attraction of pRFs. In early visual areas, where pRFs are small, position differences were largest closest to the attended locus and declined with distance from the attended locus (V1-3; Figure 3E). In higher visual areas, with larger pRFs, attraction towards the attended locus increased and plateaued with distance from the attended locus (hV4-IPS; Figure 3E). Notably, these results could only be explained by an attention field model with an offset on the attention field (AF+, Figure 4C, D, Equation 3).

While the AF model can capture the position changes in higher visual areas, it fails to capture the patterns observed in early visual cortex. The AF+ model (Figure 4C, D), which adds an offset to the attention field encompasses all the predictions of the AF model (by having an offset of zero) and further extends it (when the offset is non-zero). Notably, in our simulations we fit a single shared attention field for all visual areas field per condition. Since there are parameter trade-offs in the AF+ model, in Figure 4D, the pattern of differences in early visual areas is visible and the plateauing in higher visual areas is less well captured. This perhaps suggests that while all visual areas may share a uniform attention field size, the offset on the attention field may vary across visual areas. The AF+ model does not suffer from the limitations of the AF model and is able to capture the variety of effects observed along the eccentricity range and visual hierarchy.

Our task encouraged participants to distribute their attention in one condition by attending to a stimulus that spanned the entire screen. An inherent limitation of distributed attention paradigms is that it is possible participants differed in how much and where they allocate attention. The same limitation is present in other paradigms of distributed attention (e.g. spatial uncertainty, using a neutral cue). Even if attention was not distributed evenly across the entirety of the screen, participants are forced to distribute their attention *more* than in the focused condition. Furthermore, if attention moves to different locations in the distributed attention condition this would result in an averaged distributed attention field as we averaged all runs within each condition. So, we can deduce that even if there is variation in the degree to which participants distribute their attention this design is sufficient for our purposes: investigating the influence of precision of attention on visual responses.

Even if participants only attended to a sub-area of the distributed task stimulus, by necessity of doing the task, they would still be distributing their attention *more* than in the spatial extent of the focused attention condition (0.1° radius). Consequently, even if attention was not spread across the entirety of the screen, the distributed condition would still be more distributed than the focused condition. Furthermore, if this sub-area was consistently in one location not centered on fixation, the pattern of position changes would be towards the sub-area they were attending (Klein et al., 2014; van Es et al., 2018; Womelsdorf et al., 2006). Depending on the attended sub-area location, this would result in a well-documented but different pattern of position changes than what we observe (Klein et al., 2014; van Es et al., 2018; Womelsdorf et al., 2006). Namely, some pRFs becoming more foveal and others becoming more peripheral. Instead, we observed more foveal pRFs across the entire visual field for the focused attention condition compared to the distributed attention condition (Figure 3). If the sub-area they were attending to moved throughout the session, this would average to a distributed attention field since we averaged all runs of the same condition. In fact, this would be akin to using spatial uncertainty as a means of distributing attention (Herrmann et al., 2010; Huang et al., 2016; Itthipuripat et al., 2014). Despite being unable to resolve how much participants are distributing attention solely from behavior, the distributed attention task fulfills the intended purpose of the experiment: to focus attention in one condition and distribute it in another.

Could the observed data be explained by eye-movement differences between the conditions? We do not believe differences in eye-movements can explain our data. Unstable fixation would result in changes in pRF size but not position (Levin et al., 2010). However, in our data we observe changes in pRF position but no significant differences in pRF size (Supplementary Figure 1). Moreover, all participants were trained to fixate comparably in the two tasks outside of the scanner. Due to technical constraints, we were not able to acquire eye tracking for all subjects during scanning. For the participants for whom we also measured eye-position and movements in the scanner during the tasks. We observed no differences between conditions. Thus, differences in eye-movements can not explain our data.

Where could the attentional modulation be coming from? In the present study, we were unable to directly investigate where attentional modulation may be coming from. Based on chemical inactivation studies in primates and TMS studies in humans, frontal eye-fields, superior colliculi, and posterior parietal areas are all candidates for the source of attentional modulation (Bollimunta et al., 2018; Fernández et al., 2023; Krauzlis et al., 2013; Maunsell, 2015).

In this study we are measuring population receptive fields (pRFs). These pRFs differ from receptive fields (RFs) which are defined as stimulus locations that evoke or modulate neuronal responses (Hartline, 1938; Hubel & Wiesel, 1959; Sherrington, 1906). Population receptive fields are a statistical summary of response properties of a population, in our case a population within a voxel measured using BOLD fMRI (Dumoulin & Wandell, 2008). Previous investigations have found concordance between the properties of pRFs measured with BOLD fMRI and those measured with invasive electrophysiology (Harvey et al., 2013; Klink et al., 2021). In our data we observe a change in the position preference driven by the precision of attention. This could be due to relative amplitude changes within the population that we measure, due to a change in position preference of the underlying neuronal population, or some combination of the two. Indeed, both changes in neuronal firing rates as a result of attention (Moran & Desimone, 1985) and changes in the spatial profile of single neurons (Womelsdorf et al., 2006) have been observed using invasive electrophysiology. With the techniques used in this study, we cannot disentangle which of these phenomena is the driver of our observations.

What is the relationship between pRF position changes and attentional gain? pRF position changes and attentional gain are closely related. Gain changes at lower levels will give rise to position changes at higher levels of the hierarchy (Compte & Wang, 2006; McAdams & Maunsell, 1999; Womelsdorf et al., 2008). This relationship is captured by the attention field model: multiplying a stimulus-driven pRF by an attention field (i.e. increased gain at the attended location) produces a spatial change in the resulting attention-driven pRF (Figure 1a). In this study, we observe both gain changes as a result of the attentional task (Figure 2) and spatial position changes arising from a difference in the precision of attention (Figure 3). However, Fox and colleagues (Fox et al., 2023), using a convolutional neural network (CNN), propose that gain changes, rather than position changes, are the main driver of the behavioral enhancement of attention. By independently manipulating gain, receptive field shifts, and receptive field sizes in their CNN, they concluded that gain modulation alone accounts for most of the behavioral enhancement. In the brain, these responses do not appear in isolation, they are entangled. So, further studies are required to continue to tease apart the underlying mechanisms responsible for attentional enhancement of perception in humans.

Would our observed attentional effects alter perception? Our behavioral paradigm cannot determine whether the cortical differences due to attentional precision would also result in perceptual differences. However, previous behavioral studies indicate that attention can bias perception consistent with attraction towards the attended locus (Carrasco, 2018; Carrasco & Barbot, 2019; Klein et al., 2016).

Our results are in keeping with the ‘zoom lens’ analogy for attention (Eriksen & St James, 1986; Müller et al., 2003). In this study, we demonstrated that precision of attention altered the attraction of receptive fields towards the attended locus. Specifically, more precise, focused attention resulted in a greater attraction towards the attended locus. These position changes were observed across the entire visual hierarchy. Interestingly, we observed that this attraction also depends on the degree of overlap between a pRF and the attention field. These results were best captured by a Gaussian attention field model with an offset on the attention field.

## Supplementary Materials

**Supplementary Movie 1.**

Example video of part of the experimental stimulus and task. Pink and blue dots fill the screen, typically at a proportion of 50-50. Independently for the two conditions, there are target trials where the dot proportion deviates from 50-50 and becomes more pink or more blue. During the task a black and white checkerboard bar traverses the screen. The video is contrast enhanced for the purposes of visualization.

**Supplementary Figure 1.**
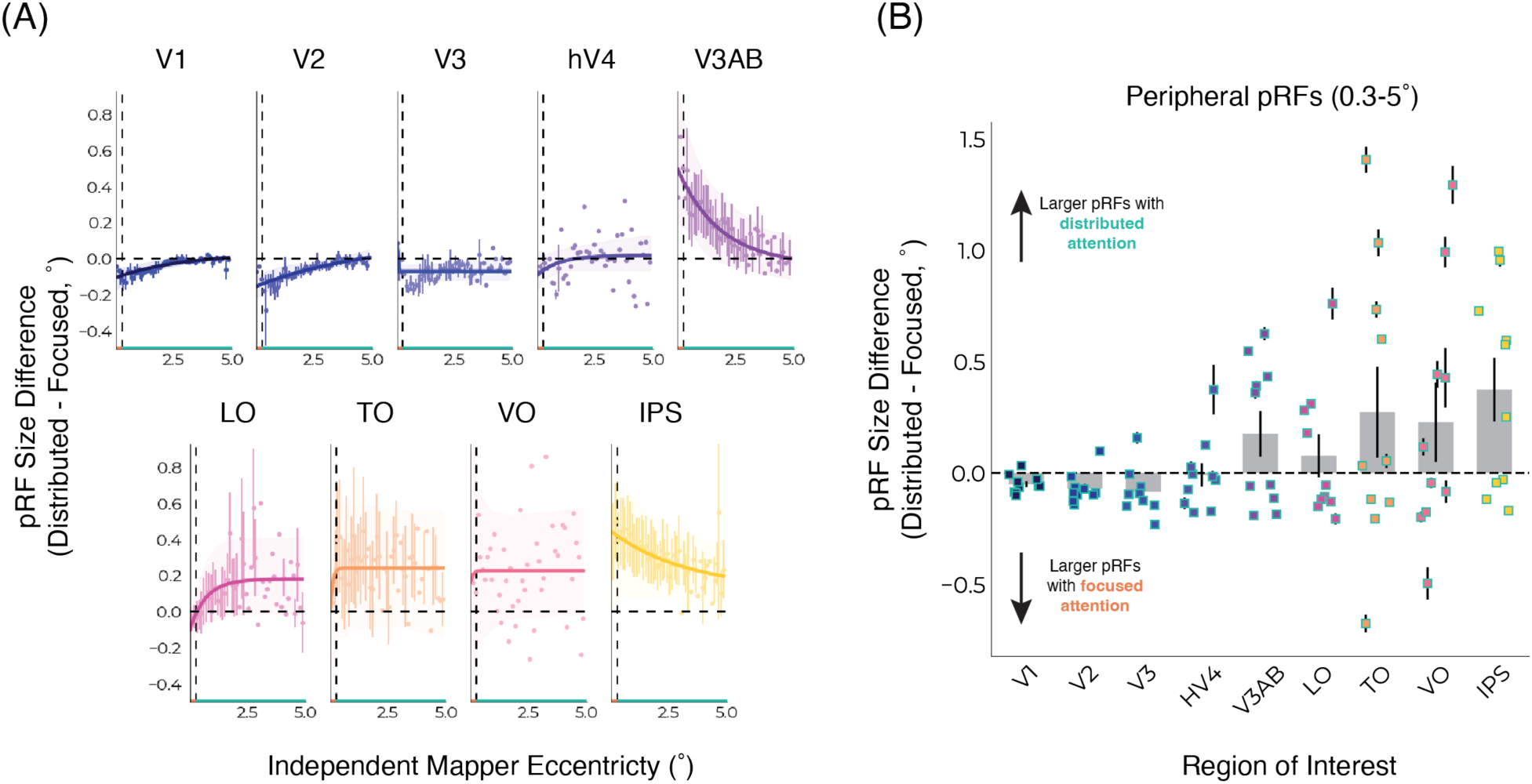
Attentional precision does not significantly alter pRF size. **a)** pRF size difference (distributed minus focused) for pRFs along the eccentricity range in each visual area. pRFs are binned for both tasks based on an independent pRF mapper. The orange line at the bottom of the plot denotes the stimulus extent of the focused attention task (0-0.3° radius), the teal line denotes the stimulus extent of the distributed attention task (0.3-5° radius), and the dashed vertical line the boundary between the two tasks. The dashed horizontal line is at 0, representing no difference between focused and distributed attention. Each individual point is an average of all participants within each eccentricity bin. The solid lines are an exponential function fit to the data. The shaded area denotes a bootstrapped 95% confidence interval of the exponential curve fit. All error bars are SEM. **b)** pRF size difference (distributed minus focused) across the visual hierarchy for pRFs falling within the distributed color task stimulus (0.3-5° radius). Values per participant are shown in the scatter plot nested within each bar, group averages in bar plots, and error bars are SEM.

