## Supplementary figures and images for "The precision of attention controls attraction of population receptive fields"

### Supplementary Movie 1

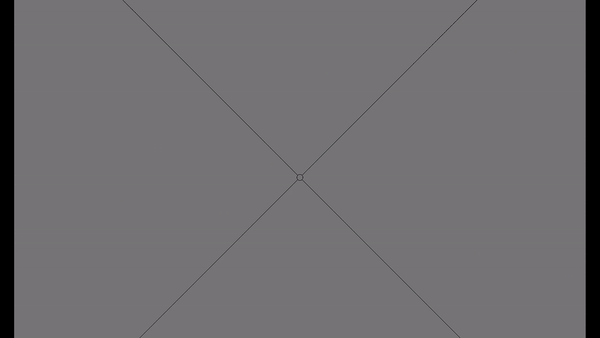
